# LOCC: a novel visualization and scoring of cutoffs for continuous variables

**DOI:** 10.1101/2023.04.11.536461

**Authors:** George Luo, John J. Letterio

**Author notes:** Corresponding Authors: George Luo, Case Western Reserve University School of Medicine, 2103 Cornell Rd., Wolstein Research Bldg. Rm 3501, Cleveland, Ohio 44106;, John J. Letterio, M.D., The Angie Fowler Adolescent and Young Adult Cancer Institute, University Hospitals Rainbow Babies & Children’s Hospital, 11100 Euclid Avenue, Cleveland, Ohio 44106.

## Abstract

**Objective:** There is a need for new methods to select and analyze cutoffs employed to define genes that are most prognostic significant and impactful. We designed LOCC (Luo’s Optimization Categorization Curve), a novel tool to visualize and score continuous variables for a dichotomous outcome.

**Methods:** To demonstrate LOCC with real world data, we analyzed TCGA hepatocellular carcinoma gene expression and patient data using LOCC. We compared LOCC visualization to receiver operating characteristic (ROC) curve for prognostic modeling to showcase its utility in understanding predictors in various TCGA datasets.

**Results:** Analysis of *E2F1* expression in hepatocellular carcinoma using LOCC demonstrated appropriate cutoff selection and validation. In addition, we compared LOCC visualization and scoring to ROC curves and c-statistics, demonstrating that LOCC better described predictors. Analysis of a previously published gene signature showed large differences in LOCC scoring, and removing the lowest scoring genes did not affect prognostic modeling of the gene signature demonstrating LOCC scoring could distinguish which predictors were most critical.

**Conclusion:** Overall, LOCC is a novel visualization tool for understanding and selecting cutoffs, particularly for gene expression analysis in cancer. The LOCC score can be used to rank genes for prognostic potential and is more suitable than ROC curves for prognostic modeling.

**Graphical Abstract:** 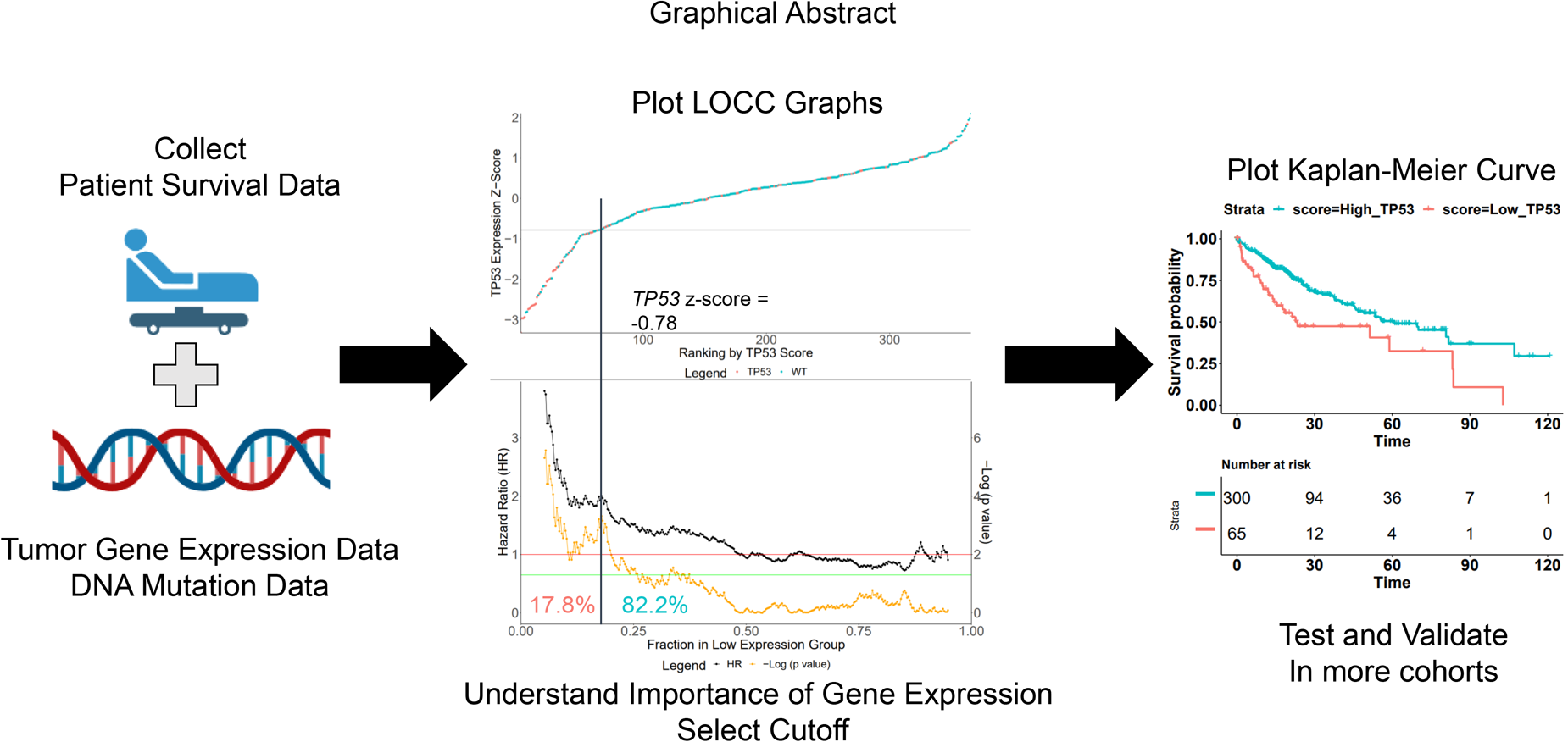

## 1. Introduction

In medicine, continuous markers refer to those that can take on a range of continuous values, rather than discrete categories. These markers can provide important information about the health status of an individual and can be used in a variety of medical applications, including diagnosis, prognosis, and treatment planning (Bennette and Vickers, 2012). Examples of continuous markers in medicine include laboratory test results such as blood glucose levels, cholesterol levels, and white blood cell count. For cancer, examples include tumor size, gene expression, and many other biomarkers such as prostate specific antigen (Bennette and Vickers, 2012; Eng, et al., 2015). Use of these markers can be helpful for both the patient and physician.

The interpretation of continuous markers can also be challenging, and this is especially true for new biomarkers that are just being tested. The most important question for new biomarkers is where to draw the line to divide groups (Altman and Royston, 2006; Budczies, et al., 2012). While traditional methods have used median or quartiles, more recent computational methods choose the cutoff that has the most significant p-value (Bennette and Vickers, 2012; Budczies, et al., 2012). However, in both cases, there is a significant loss of information as all other possible cutoffs are ignored and only information using the specific cutoff is relayed to the user. There are other computational tools which also try to convey information about the p-value or hazard ratio over the range of the continuous variable yet we have not found a comprehensive method that allows visualization of the full picture (Budczies, et al., 2012; Nagy, et al., 2021).

Current calculation methods employed to evaluate models for prognosis and outcome are also challenged by limitations. A popular method is use of receiver operating characteristic (ROC) curve, a plot of sensitivity and specificity (Cook, 2008; Zou, et al., 2007). The area under the curve (AUC) provides information whether the variable has good performance at distinguishing the outcome. While ROC is appropriate for definitive diagnosis, it lacks discrimination and calibration in most prognostic studies (Berrar and Flach, 2012; Cook, 2008; Grund and Sabin, 2010; Zou, et al., 2007). For example, a PCR test for HIV would be appropriate for ROC evaluation but single gene expression in cancer biopsy is too little information to determine the consequences. Hence, there is a need for better calculations for continuous variables relating to outcomes.

Therefore, we developed Luo’s Optimization Categorization Curves (LOCC) to help visualize more information for better cutoff selection and understanding of the importance of the continuous variable against the measured outcome. We describe the process of understanding and generating LOCC with practical survival curve examples using real data from The Cancer Genome Atlas (TCGA, 2017) while comparing it to ROC curve analysis. We also reanalyzed published data and optimize gene signatures using a LOCC score to evaluate the gene expression significance and impact on prognosis. We believe LOCC will lead to improved analysis and understanding of continuous variables in many biological settings.

## 2. Methods

### 2.1 Data Sources

We used TCGA data from cBioportal.com (Cerami, et al., 2012) and LIRI-JP (Liver hepatocellular carcinoma – Japan) data from International Cancer Genome Consortium (ICGC). Z-scores of gene expression analyzed by RNA-Seq by Expected Maximization (RSEM) for TCGA was used while RNA-sequencing data was acquired in normalized read counts for LIRI-JP data. Normalization of read counts to z-scores was performed for each sample by subtracting the individual expression by the mean expression and dividing by standard deviation. Mutation data was acquired for TCGA data and considered mutant for any non-conservative mutation.

### 2.2 LOCC Visualization

To generate the LOCC ranking graphs, data was processed in R. Samples were ordered by expression z-scores and then graphed using ggplot. A horizontal line representing the ideal or selected cutoff was added as needed.

To generate the LOCC cutoff selection, we analyzed possible categorization by analyzing every cutoff according to the gene expression that would result in a different grouping of patient samples and calculating the corresponding hazard ratio (HR) and p-values. R package survival was used to calculate the hazard ratios and p-values. The HR was calculated with a cox proportional hazard regression model. A log-rank test was used to evaluate p-values. For n number of patients, we would perform n-1 number of cuts to calculate corresponding HR and p-values. We graph the HR and p-values for each cutoff that has a minimum of 5% of the total population in each group. For the optimal cutoff, the cutoff with the lowest p-value in which each group has at least 10% of the population is selected. The cutoff was confirmed with cutpointR (Budczies, et al., 2012). Code for LOCC algorithm will be available on publication.

### 2.3 LOCC Score

The LOCC score is made of three numeric components: a significance aspect, a range aspect, and an impact aspect. The significance is represented by the −log (p-value). The range is represented by the percentage of cutoffs that has a p-value below

0.01. The impact is represented by the highest HR, which includes ranking expression from highest to lowest as well as lowest to highest. The three numbers are multiplied together for the LOCC score, represented by the following equation.

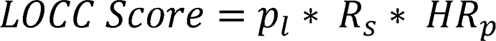

Where *p_l_* is the highest value of −log (p value), *R_s_* is percentage of cutoffs that have a highly significant p value (p < 0.01), and *HR_p_* is the HR at the most significant point. For significance and impact, the numbers are restricted to cutoffs such that at least 10% of the population is in each group. For the LOCC score, variables can be ordered by lowest to highest or vice versa such that the HR is above one.

### 2.4 ROC Curve

After processing the patient and tumor data, ROC curves were generated in R using the ROCR package. The AUC was calculated with ROCR using overall survival while only removing patients with incomplete survival or expression data. A red line of sensitivity = (1-specificity) was added for comparison.

### 2.5 RISK Gene Signature

The risk score analysis for hepatocellular was used from a previous publication (Ouyang, et al., 2020) using the same gene expression and weight coefficients. Cox regression analysis was calculated with Cox proportional hazard package in R. ROC analysis was conducted with ROCR package. ROC calculations were performed after selecting patients who survived at least 1 month. Gene expression correlations were performed from cBioportal.com (Cerami, et al., 2012).

## 3. Results

### 3.1 LOCC demonstrates *E2F1* expression is associated with a poor prognosis in hepatocellular carcinoma

To demonstrate the utility of LOCC, we used TCGA data to showcase how LOCC can help analyze the role of transcription factor *E2F1* in liver hepatocellular carcinoma (TCGA LIHC). *E2F1* is an important transcription factor that has roles in cell cycle, DNA repair, and even apoptosis (Kent and Leone, 2019; Meng and Ghosh, 2014). *E2F1* can bind p53 to induce apoptosis and can also be inhibited by the retinoblastoma protein (Rb) to arrest the cell at the G1/S checkpoint (Kent and Leone, 2019; Meng and Ghosh, 2014). As such, *E2F1* is an important target in cancer where it is often overexpressed.

LOCC is a comprehensive visualization of multiple parameters for continuous variables (See Supplemental Fig. 1 for detailed labeling of LOCC). The first graph, the LOCC ranking, plots the values of the continuous variable against the ranking of all samples. In our case, we plot *E2F1* z-score expression in the tumor samples on the y-axis while we show the ranking of the samples on the x-axis (Fig. 1A). We use the z-score because many datasets have different methods of standardizing RNA-Seq data, but we use a normalization method to approximate the distribution. We also plot mutations of this gene with different colors to see if mutations affect gene expression. Next, we graph the LOCC cutoff selection, which plots the hazard ratio at every single cutoff (Fig. 1B). We observe the hazard ratio (black line) is almost always above 1 (red line) which means that a higher *E2F1* expression is almost always associated with a worse prognosis in LIHC.

**Figure 1.**
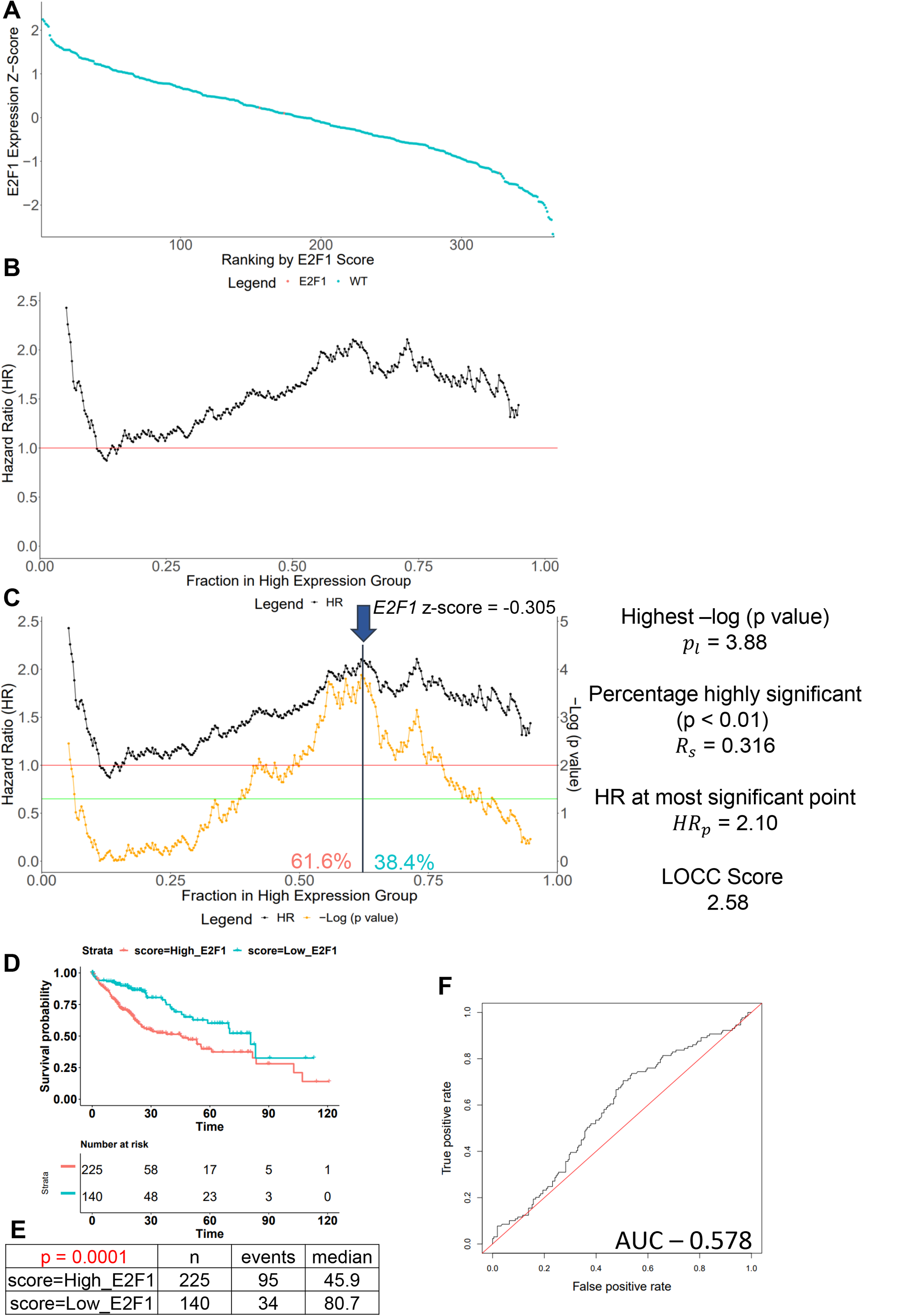
LOCC demonstrates *E2F1* is associated with a poor prognosis in TCGA hepatocellular carcinoma. **(A)** The z-score expression of *E2F1* from TCGA hepatocellular carcinoma patient samples was ordered in descending order and plotted against the ranking of the samples. Samples with non-conservative mutations of *E2F1* are colored in orange while samples with wildtype *E2F1* are colored turquoise. **(B)** A black line depicting the hazard ratio is plotted for every cutoff for *E2F1* expression. A red horizontal line is place at HR = 1.0. (**C**) A yellow line depicting the −log (p-value) is added to the graph to display the significance of each cutoff. The red horizontal line is also aligned with p = 0.01 while the green horizontal line is aligned with p = 0.05. The cutoff with the lowest p-value is selected to be the ideal cutoff, indicated by the arrow which corresponds to a z-score of −0.305. (**D**) A Kaplan-Meier overall survival curve is plotted at the ideal cutoff to separate patients into high or low *E2F1*. A data table of the details of the groups is shown. Patients’ survival times are expressed in months. P-values are calculated using log-rank test. HR are calculated using Cox proportional hazard regression.

However, an important question is whether *E2F1* expression is truly significant at any point. To answer this question, we add a second line defined by its own y-axis scale, −log (p-value), on the hazard ratio graph (Fig. 1C). The key values for the yellow – log (p-value) line is p = 0.01 (red horizontal line) and p = 0.05 (green horizontal line). Therefore, whenever the yellow line is above the green line, it is statistically significant (p < 0.05) and above the red line is very significant (p < 0.01). In this case, the ideal cutoff should be between the top 50% and top 75% of *E2F1* expression. Using a lowest p-value approach, a cutoff at 61.6% yields the lowest p value and seems appropriate. At the same time, the exact *E2F1* expression at this cutoff can be visualized using the first graph shown in Fig. 1C. We also create the Kaplan-Meier curve at that cutoff to see its appropriateness and calculate a median survival (Fig. 1D-E). Previous literature has demonstrated *E2F1* gene expression is a prognostic marker for hepatocellular carcinoma using a median expression cutoff (Huang, et al., 2019).

We have also developed a LOCC score to help judge significance and overall impact of any single or group of predictors. Three factors are multiplied together to get the LOCC score: the significance, the range, and the impact. The significance is calculated by taking the −log (lowest p-value). The range is the percentage of LOCC significance line (yellow line) that is above the red line (p < 0.01). Finally, the impact is the HR at the most significant cutoff (if HR is below 1, use the reciporal). We also limit group sizes for significance and impact to be at least 10% of the population to minimize extreme effects from small samples. Thus, the higher the LOCC score, the more critical and predictive the expression is for prognosis. In our evaluation of *E2F1,* the LOCC score was calculated to be 2.58 (Fig 1C), which demonstrate there is some prognostic value in this dataset. In the following sections, the scale of the LOCC score will be established.

We also generated a ROC curve using *E2F1* expression and survival status (Fig. 1F). The area under the curve (AUC) was 0.578 which suggested there was increased risk of death with higher *E2F1* expression. Interestingly, we noticed that the red line for LOCC (HR = 1) and ROC (True Positive Rate = False Positive Rate) were actually the same underlying equation (Supplemental Figure 2A-E). If the HR line is above the red line for the entirety of the graph, then the same is true of the ROC curve above the red line of random classifier. Therefore, the relative positions (above/below) of the black and red lines should be similar in both LOCC and ROC curves.

### 3.2 Validation of *E2F1* as prognostic biomarker in the LIRI-JP hepatocellular dataset

After finding a very significant cutoff and setting the cutoff, we need to validate this result in another cohort of hepatocellular carcinoma. For this, we used data from the International Cancer Genome Consortium (ICGC) which had a large Japanese cohort of liver hepatocellular carcinoma (LIRI-JP). The LIRI-JP data gives normalized read counts represented by Fragments Per Kilobase of transcript per Million mapped reads (FPKM), which is different from the RNA-Seq by expectation maximization (RSEM) used by TCGA. While it is possible to reprocess the raw data of both groups with the same programs to end up with the same expression format, that process would require tremendous effort. Instead, for this example, we will use the normalized z-score to estimate the cutoff which should approximately judge the relative expression between patients, though it is imperfect. While the z-scores range and distribution may vary significantly between populations, we hope to capture a similar categorization of the population in which we believe there are at least two groups that can be stratified by *E2F1* expression.

We used the *E2F1* cutoff from the TCGA training of −0.305 z-score and set that as the cutoff for the validation cohort (Fig. 2A). Using LOCC, we can analyze how significant and appropriate that cutoff is (Fig. 2B). Even though the cutoff is not quite the highest peak of significance, it is still highly significant so the cutoff is still adequate. Using the cutoff, we generate a Kaplan-Meier plot to visualize the survival curves between the two groups (Fig. 2C). With significant survival differences in both TCGA and LIRI-JP cohorts using the same cutoff (Fig. 2D), it would be appropriate to suggest that *E2F1* is a prognostic biomarker for LIHC associated with a poor prognosis. The LOCC score was calculated to be 18.9 (Fig. 2B), which suggests that *E2F1* is of great prognostic potential in this dataset.

**Figure 2.**
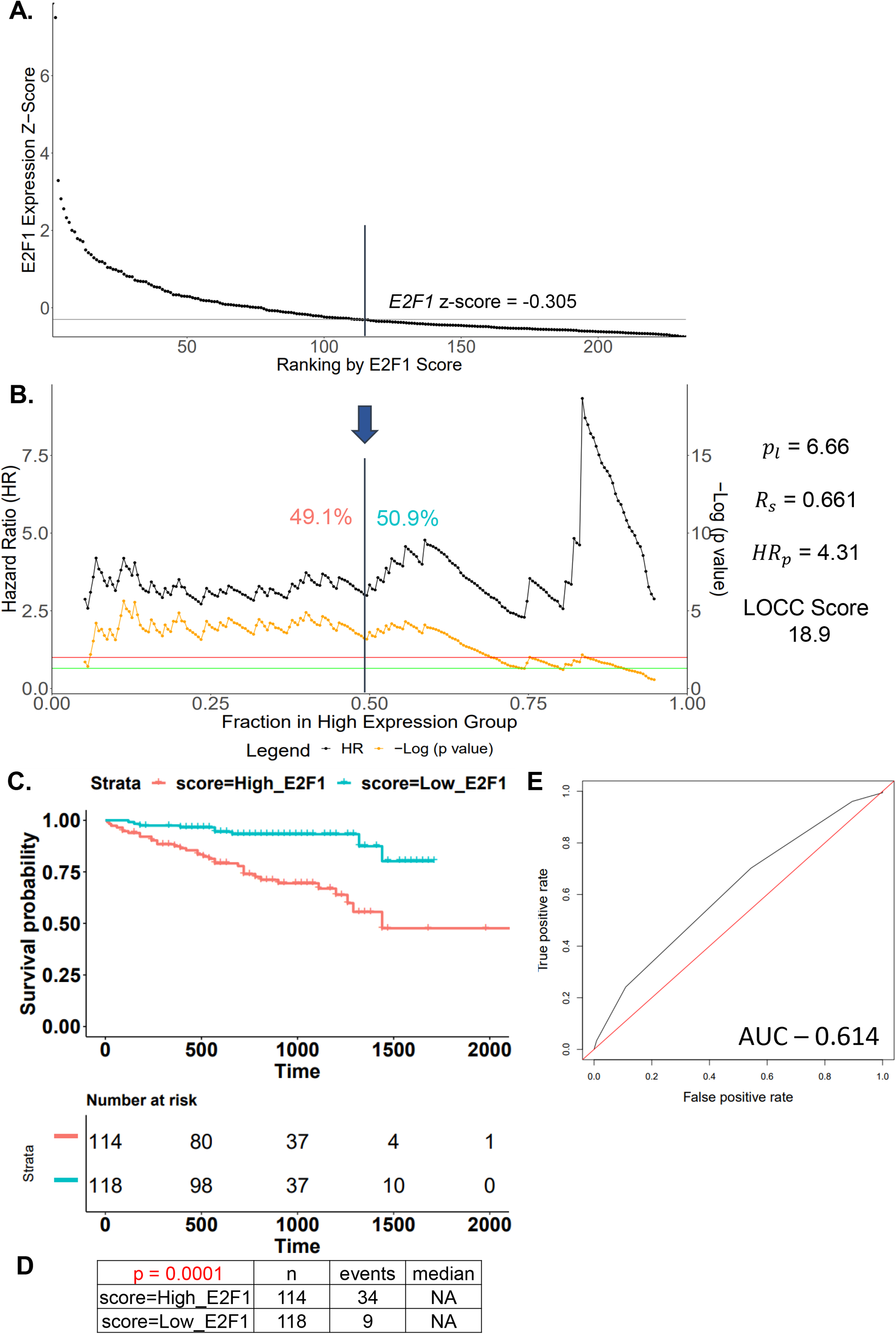
Validation of *E2F1* as prognostic biomarker in the LIRI-JP hepatocellular dataset. **(A)** The z-score expression of *E2F1* from LIRI-JP hepatocellular carcinoma patient samples was ordered in descending order and plotted against the ranking of the samples. A horizontal line is graphed at −0.305 to separate patients into high and low *E2F1* groups using the TCGA dataset cutoff. **(B)** The LOCC cutoff selection was graphed for LIRI-JP samples. The cutoff from TCGA data was used to separate patients into high and low *E2F1* groups. (**C**) A Kaplan-Meier overall survival curve is plotted at the validation cutoff to separate patients into high or low *E2F1*. A data table of the details of the groups is shown. Patients’ survival times are expressed in days. P-values are calculated using log-rank test. HR are calculated using cox proportional hazard regression.

With the ROC curve, we can see that the AUC is 0.614, suggesting there is increased risk of death associated with *E2F1* expression. However, it is not easy to see how significant this predictor is or what cutoffs are optimal or significant. As such, the evaluation of *E2F1* expression as a prognostic biomarker by ROC is minimal compared to LOCC.

### 3.3 Comparison of ROC and LOCC in evaluating prognostic biomarkers

ROC curve and the AUC is commonly used in many biomarkers and prognostic modeling (Cook, 2008; Zou, et al., 2007). However, the interpretation of the ROC graph and the AUC (also known as the c-statistic) is questionable for prognosis. For diagnosis, the c statistic can be thought of as the probability in which a randomly selected subject who experienced the event had a higher test score than a subject who did not experience the event (Austin and Steyerberg, 2012). The c-statistic, in the case of prognosis, is the probability of a randomly selected patient who experienced death had a higher test score than a patient who did not die. While c-statistics is appropriate for diagnosis, its meaning is difficult to interpret for prognosis. Simply classifying patients as alive or dead is not sufficient for overall survival and inappropriate for censored data. Even though time-dependent ROC attempts to resolve the first issue, it still struggles with censorship (Kamarudin, et al., 2017).

Instead, with LOCC, we use the cox regression model and investigate hazard ratios between groups. By classifying patients into two groups based on the biomarker, we can evaluate the hazard ratio and p values. For LOCC, we do this for all possible cutoffs, similarly to ROC, and plot the hazard ratio and p values to find the best cutoffs. This not only helps with selection of the cutoff but also allows us to evaluate the changes in hazard ratio over the range of the biomarker.

In the TCGA Sarcoma (SARC) dataset (TCGA, 2017), a ROC curve is plotted for *E2F1* expression where the AUC is 0.576 (Fig. 3A). Despite this AUC being similar to the AUC for *E2F1* expression in LIHC, the LOCC graph looks very different and the LOCC score for sarcoma *E2F1* is only 0.11 (Fig. 3B), much lower than the 2.58 for *E2F1* in LIHC. The primary difference between the LOCC scores is the p values and highly significant ranges while the HR were similar. After selecting the most significant cutoff, a Kaplan-Meier curve is plotted (Fig. 3C). However, with a low LOCC score, our interpretations is that the prognostic value of *E2F1* in sarcoma is not as robust as *E2F1* in hepatocellular carcinoma.

**Figure 3.**
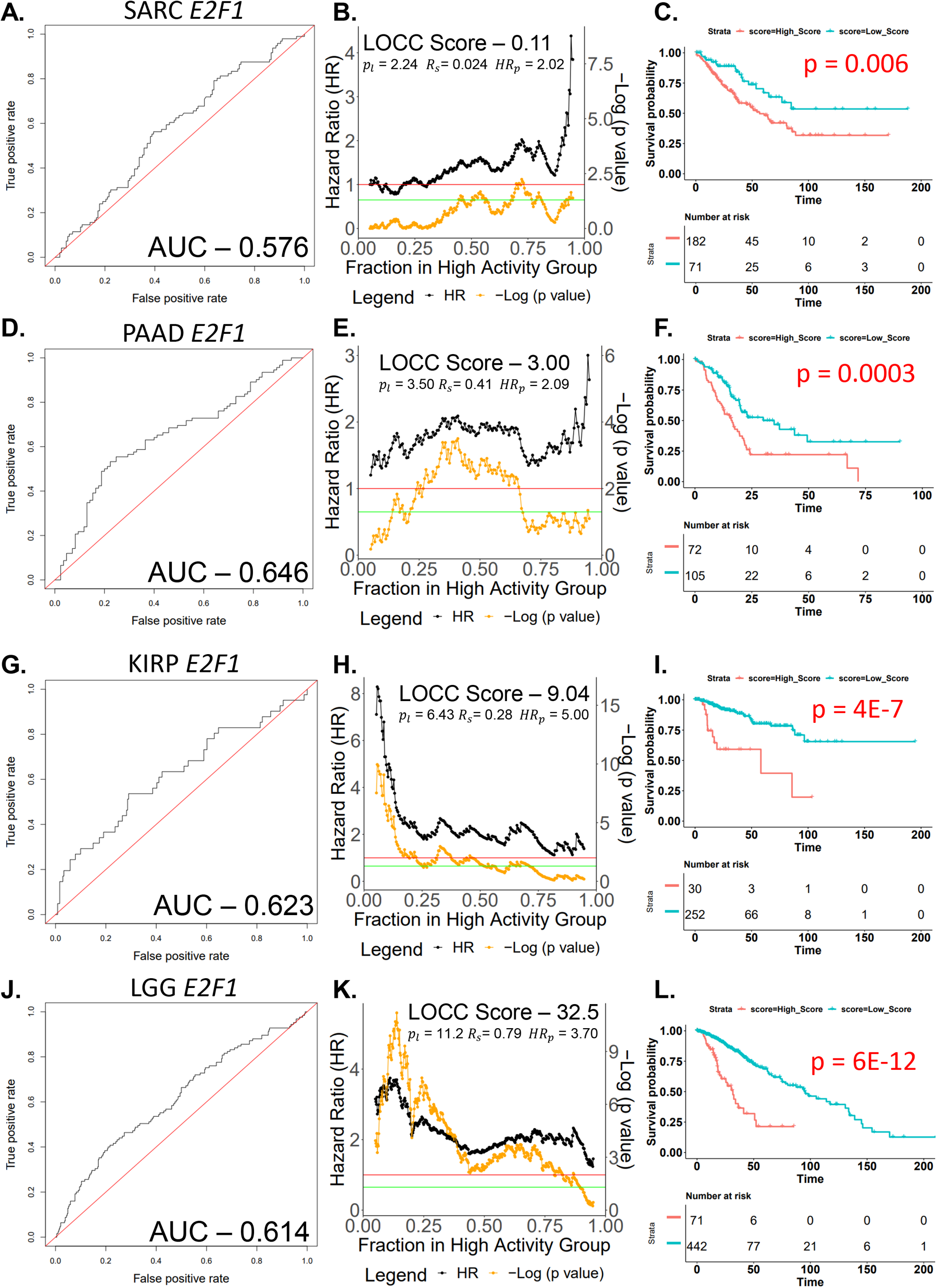
Comparison of ROC and LOCC in evaluating prognostic biomarkers. **(A)** A ROC curve is plotted and the AUC was calculated for TCGA SARC *E2F1*. (**B**) LOCC was plotted and scored for SARC *E2F1.* (**C**) The most significant cutoff was selected and a Kaplan-Meier plot was graphed to illustrate the best stratification according to *E2F1* in SARC. **(D)** A ROC curve is plotted and the AUC was calculated for TCGA PAAD *E2F1*. (**E**) LOCC was plotted and scored for PAAD *E2F1.* (**F**) The most significant cutoff was selected and a Kaplan-Meier plot was graphed to illustrate the best stratification according to *E2F1* in PAAD. **(G)** A ROC curve is plotted and the AUC was calculated for TCGA KIRP *E2F1*. (**H**) LOCC was plotted and scored for KIRP *E2F1.* (**I**) The most significant cutoff was selected and a Kaplan-Meier plot was graphed to illustrate the best stratification according to *E2F1* in KIRP. **(J)** A ROC curve is plotted and the AUC was calculated for TCGA LGG *E2F1*. (**K**) LOCC was plotted and scored for LGG *E2F1.* (**L**) The most significant cutoff was selected and a Kaplan-Meier plot was graphed to illustrate the best stratification according to *E2F1* in LGG. Abbreviations and symbols: TCGA – The Cancer Genome Atlas, SARC – Sarcoma, PAAD – Pancreatic ductal adenocarcinoma, KIRP – Kidney renal papillary cell carcinoma, LGG – low grade glioma, *p_l_* is −log (p value) at most significant cutoff, *R_s_* is percentage of cutoffs that have a highly significant p value (p < 0.01), *HR_p_* is the HR at the most significant cutoff.

In the TCGA pancreatic ductal adenocarcinoma (PAAD) dataset (TCGA, 2017), we plotted the ROC curve for *E2F1* and observed an AUC of 0.646 (Fig. 3D). This is a relatively high AUC and our LOCC score is 3.00 (Fig. 3E), an increase from SARC data just like the AUC. Yet, we can also observe that the HR between the two groups is about 2.0 for both cancers at the most significant cutoffs. However, the p values throughout the LOCCs are different, leading to the discrepancy in scores. Using the most significant cutoff, a Kaplan-Meier plot was generated which demonstrates a significant stratification using *E2F1* expression (Fig. 3F).

In the TCGA kidney renal papillary cancer (KIRP) dataset (Linehan, et al., 2016), we plotted the ROC curve for *E2F1* and observed an AUC of 0.623 (Fig. 3G). On initial inspection, the KIRP ROC curve looks similar to the PAAD curve. However, LOCC paints a different picture with a very high-risk group among the top 20% of *E2F1* expression while the differences using cutoffs near the median are borderline significant (Fig. 3H). The LOCC score of 9.04 also demonstrate there is high prognostic potential of *E2F1* in this dataset if the correct cutoff is used. The Kaplan-Meier curve reflects that a small proportion of KIRP patients are at significantly higher risk with high *E2F1* expression (Fig. 3I).

Finally, using TCGA lower grade glioma (LGG) data (Brat, et al., 2015), we plotted the ROC curve for *E2F1* and found an AUC of 0.614 (Fig. 3J). Despite having a lower AUC compared to the AUC from PAAD and KIRP, LGG LOCC score of 32.5 was much higher and showed most cutoffs were highly significant (Fig. 3K). This can be also seen in the Kaplan-Meier curve where two very different groups of patients can be distinguished using *E2F1* (Fig. 3L). Therefore, *E2F1* expression is a very prognostic significant predictor in LGG and other researchers have investigated targeting this pathway (Wang, et al., 2021).

### 3.4 Low expression and mutations of *TP53* are associated with a poor prognosis

Next, we demonstrate LOCC utility in helping evaluate gene expression in genes with prognostic significant mutations. For TCGA LIHC, *TP53* mutation is prognostic significant for survival as patients with mutations have a worse prognosis (p – 0.04, Fig. 4A). However, there are other mechanisms leading to disruption of the p53 pathway besides mutations and LOCC can help evaluate them. Low *TP53* expression is also significantly associated with a poor prognosis (p – 0.0005, Fig. 4B). A z-score *TP53* expression plot demonstrates that *TP53* mutations are associated with low expression of *TP53* (Fig. 4C). We also ranked *TP53* expression from low to high so the LOCC graph will show a significant HR above 1, reducing overlapping lines and making it easier to read the graph (Fig. 4D). Therefore, both expression and mutation of TP53 can be used in combination to help determine the prognosis of patients with LIHC, which is in line with previous studies (Donehower, et al., 2019; Yang, et al., 2022).

**Figure 4.**
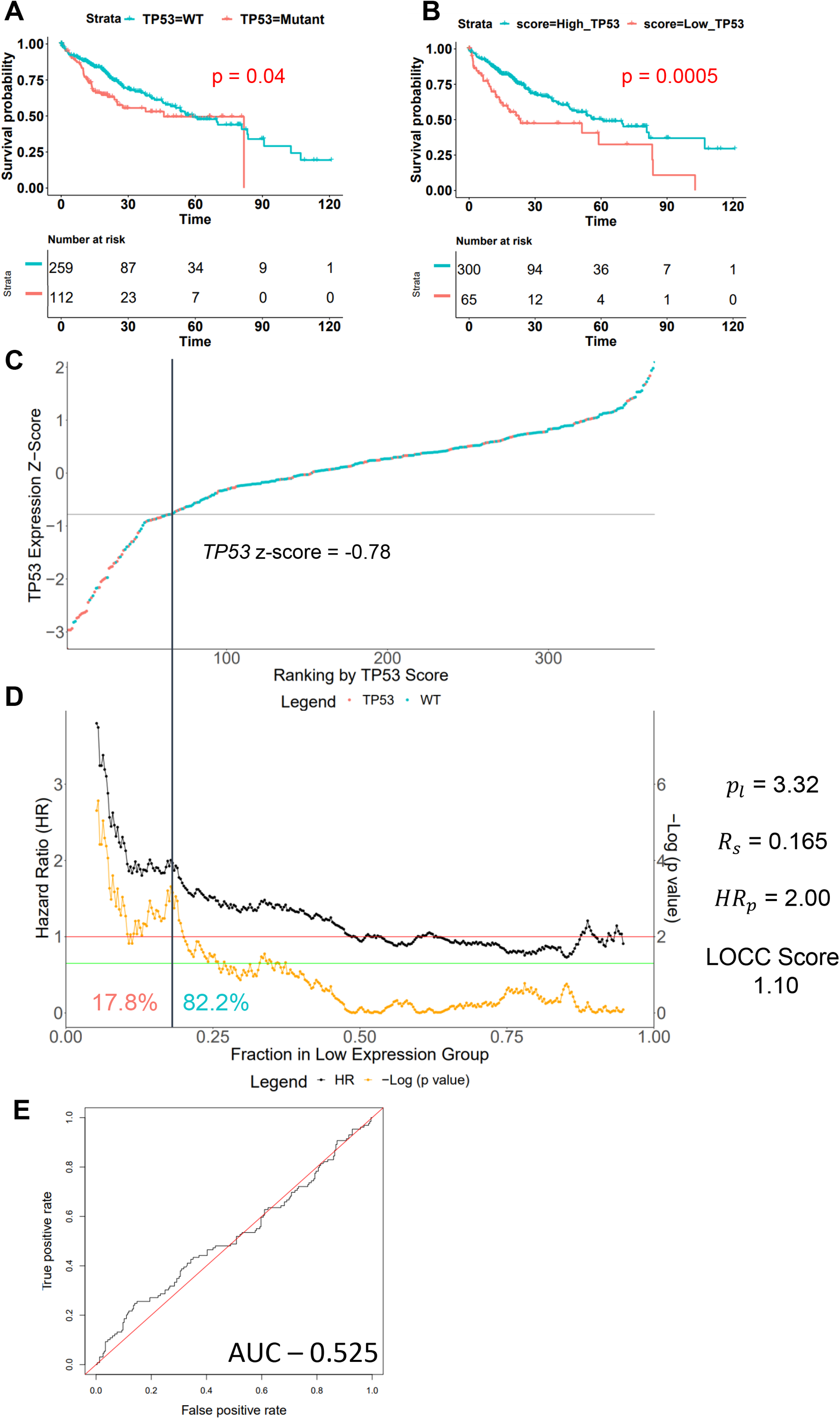
Low expression and mutations of *TP53* are associated with a poor prognosis in TCGA hepatocellular carcinoma. **(A)** The Kaplan-Meier survival curve is plotted for *TP53* mutations. **(B)** The Kaplan-Meier survival curve is plotted for *TP53* expression with groups determined by the ideal cutoff in LOCC visualization. (**C**) The LOCC ranking is graphed for *TP53* expression z-score from lowest expression to highest expression. Samples with *TP53* non-conservative mutations are colored in orange while samples with wildtype *TP53* are colored in turquoise. (**D**) The LOCC cutoff selection is graphed for TCGA hepatocellular carcinoma *TP53* expression. The ideal cutoff is selected and this corresponds to the *TP53* z-score of −0.78. Patients’ survival times are expressed in days. P-values are calculated using log-rank test. HR are calculated using Cox proportional hazard regression.

### 3.5 LOCC score helps rank prognostic importance of predictors

We used LOCC scoring to rank the 12 individual genes from a previously published prognostic gene signature (RISK) in hepatocellular carcinoma (Ouyang, et al., 2020). We found the top eight gene expression by LOCC score (*KIF20A, TTK, TPX2, LCAT, SPP1, HMMR, CYP2C9, ANXA10*) had a significantly higher LOCC score than the other 4 genes (log_10_ LOCC p-value < 0.01, Table 1). This was different from ROC analysis in the article where the AUC of the 12 genes were all between 0.57 and 0.63 and 10 genes between 0.60 and 0.63 (Ouyang, et al., 2020). Meanwhile, cox regression analysis also found a significant difference in the −log (p-value) between the 8 genes from LOCC analysis and the other 4 genes (p = 0.0007, Table 1). We tested if this new gene signature (8-gene RISK) would be as useful as the original 12 as it contained the most significant genes according to LOCC scoring. To do this, we used the same weight coefficient as the original article and used it to calculate the 8-gene RISK score (Ouyang, et al., 2020). Indeed, we found that the optimized gene signature had a similar p-value and HR between the high and low risk groups compared to the original 12 gene signature (Fig 4A-D, Supplemental Figure 3A-C). Furthermore, the LOCC score, AUC of the ROC curves, and cox regression analysis of the 8-gene RISK and original 12-gene RISK produced very similar numbers. (Fig 4D and Supplemental Fig 3D-G). Overall, from the extensive analysis and comparisons, we believe the gene signature from the top 8 genes by LOCC score is non-inferior to the original 12 gene original signature demonstrating LOCC score ability to decipher the key predictors.

**Figure 5.**
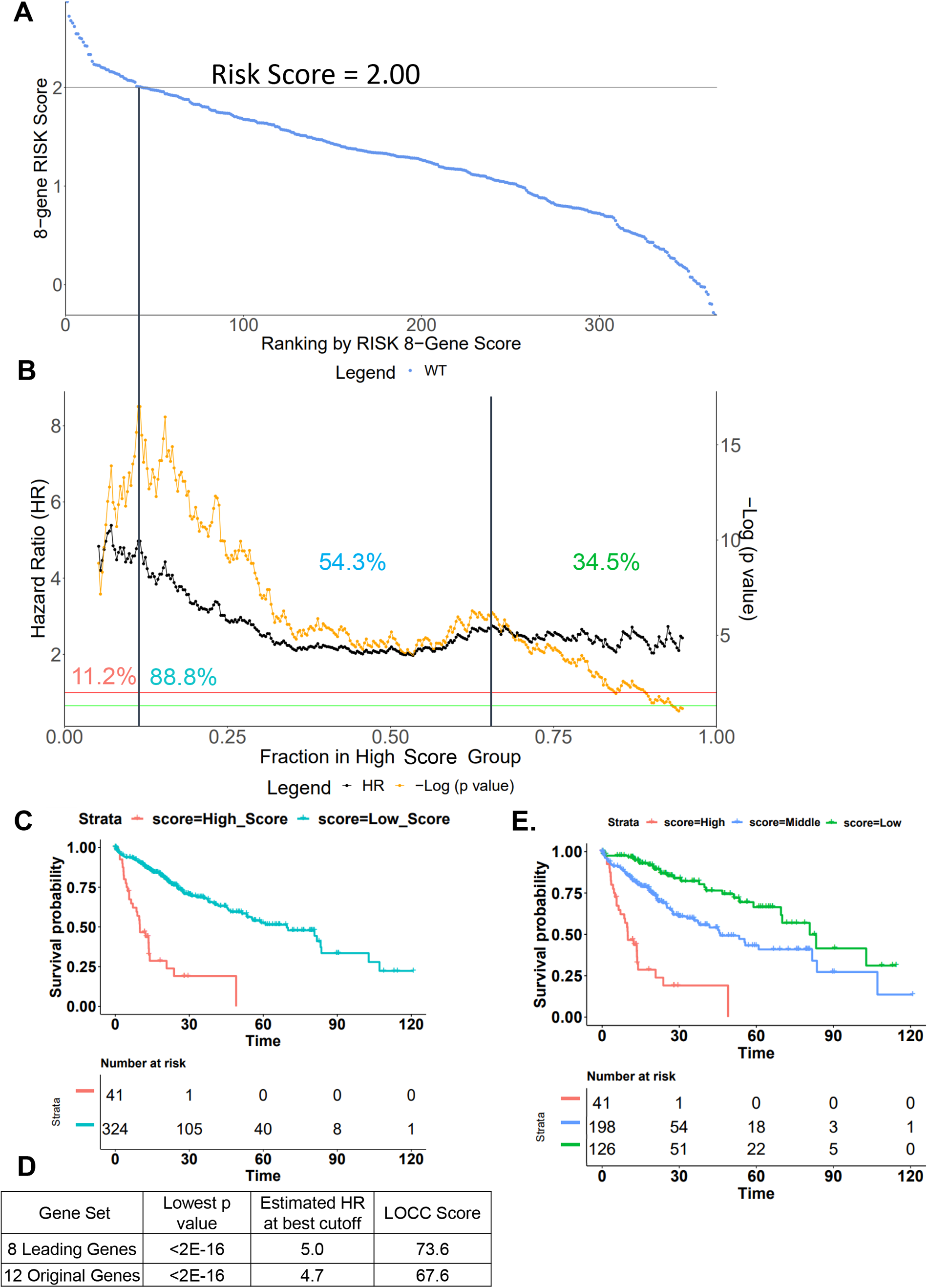
LOCC score helps rank prognostic importance of predictors. **(A)** The 8-gene modified RISK score was ordered in descending order and plotted against the ranking of the samples. A horizontal line is graphed at 2.00 to separate patients into high and low risk groups using ideal LOCC cutoff. **(B)** The LOCC cutoff selection was graphed for the 8-gene RISK score using TCGA hepatocellular data. The most significant cutoff was chosen to separate patients into two groups. In addition, another cutoff at 1.077 was selected for dividing patients into three groups. (**C**) A Kaplan-Meier overall survival curve is plotted at the validation cutoff to separate patients into high or low risk score. (**D**) A comparison of the RISK score of 8 leading genes by LOCC score and the original 12 gene RISK score is shown. The lowest p-value and hazard ratio (HR) are selected with a minimum of at least 10% of the total samples in each group. (**E**) Low risk patients are further stratified using the LOCC cutoff selector into middle and low risk. A corresponding Kaplan-Meier graph is plotted for the three different risk group. Patients’ survival times are expressed in months. P-values are calculated using log-rank test. HR are calculated using Cox proportional hazard regression.

**Table 1.** LOCC score of RISK gene signature. A table showing the analysis of the 12 hepatocellular carcinoma gene signature using LOCC score. The bolded genes are ones used for the 8-gene risk gene signature.

| Gene | -log (lowest p value) | Highly significantly Range (p <0.01) | Highest HR | Lowest HR | HR at Significant Cutoff | LOCC Score | Log<sub>2</sub> LOCC Score | Good or Poor Prognosis | Cox Regression exp(β) | Absolute Value (β) | Cox Regression p value | -log(cox p value) |
| --- | --- | --- | --- | --- | --- | --- | --- | --- | --- | --- | --- | --- |
| KIF20A | 16.097 | 0.709 | 4.094 | 1.470 | 4.094 | 46.7 | 5.545 | Poor | 1.320 | 0.278 | 0.000006 | 5.22 |
| TTK | 9.874 | 0.694 | 3.291 | 1.232 | 3.291 | 22.1 | 4.466 | Poor | 1.260 | 0.231 | 0.00002 | 4.70 |
| TPX2 | 5.650 | 0.818 | 2.761 | 1.661 | 2.761 | 12.7 | 3.667 | Poor | 1.375 | 0.318 | 0.000004 | 5.40 |
| LCAT | 8.077 | 0.598 | 0.714 | 0.381 | 0.381 | 12.6 | 3.655 | Good | 0.795 | 0.229 | 0.000001 | 6.00 |
| SPP1 | 5.692 | 0.799 | 2.415 | 1.740 | 2.312 | 10.5 | 3.392 | Poor | 1.125 | 0.118 | 0.000004 | 5.40 |
| HMMR | 5.363 | 0.780 | 2.480 | 1.417 | 2.199 | 9.2 | 3.202 | Poor | 1.299 | 0.262 | 0.00002 | 4.70 |
| CYP2C9 | 5.599 | 0.658 | 0.677 | 0.410 | 0.449 | 8.2 | 3.036 | Good | 0.900 | 0.105 | 0.0001 | 4.00 |
| ANXA10 | 4.993 | 0.738 | 0.781 | 0.420 | 0.469 | 7.9 | 2.982 | Good | 0.887 | 0.120 | 0.0001 | 4.00 |
| RDH16 | 5.332 | 0.562 | 0.689 | 0.452 | 0.452 | 6.6 | 2.722 | Good | 0.897 | 0.109 | 0.0002 | 3.70 |
| LAPTM4B | 4.836 | 0.543 | 2.532 | 1.444 | 2.094 | 5.5 | 2.459 | Poor | 1.270 | 0.239 | 0.0003 | 3.52 |
| LECT2 | 5.041 | 0.501 | 0.847 | 0.430 | 0.430 | 5.9 | 2.555 | Good | 0.920 | 0.083 | 0.0002 | 3.70 |
| MAGEA6 | 3.622 | 0.394 | 2.518 | 0.874 | 2.335 | 3.3 | 1.736 | Poor | 1.060 | 0.058 | 0.001 | 3.00 |
| Top 8 vs Bot 4 (t test p value) | 0.066 | 0.003 | 0.482 | 0.385 | 0.380 | 0.051 | 0.004 |  | 0.499 | 0.135 | 0.134 | 0.00070 |

Finally, we show that it is possible to use multiple cutoffs if there are multiple peaks on the yellow line above the red line. We chose two peaks on the LOCC cutoff selection and as such, separated patients into three well-stratified groups of high, middle, and low risk (Fig 4B and E). As such, with LOCC, it can be quickly determined whether multiple stratifications are appropriate and possible with the predictor and data.

## 4. Discussion

In this study, we demonstrate a novel visualization tool, LOCC, that presents the distribution, significance, and hazard ratios related to a continuous variable. Using this information, either an optimized cutoff or multiple cutoffs can be chosen while seeing the whole distribution. There are a number of advantages to LOCC over previous methods for prognosis. By combining hazard ratio and p-value into a single graph, one can simultaneously understand both the significance and effect of a variable. The LOCC ranking can show the distribution and mutation status of the samples. These characteristics of LOCC help analysis of the variable in relationship to the outcome and mutation status such as the case with *TP53*. By directly comparing ROC curves and LOCC, we demonstrate that there is much more helpful information present in LOCC compared to ROC curves. Sensitivity, specificity, and the c-statistic in the settings of prognosis is not easy to interpret while HR and p values are simple to understand. Therefore, we believe this LOCC visualization tool can be useful to many continuous variables to outcome analyses, and outperform ROC curves (Grund and Sabin, 2010).

We proposed a LOCC score that is an interesting metric for ranking variables by significance, range, and impact. Using a combination of three variables, we can generate a score to help determine the most valuable variables related to the outcome. The LOCC score appears to be better for prognostic studies than ROC as the AUC cannot discriminate between variables if only modest differences exists. Additionally, LOCC incorporates significance into its scoring while ROC does not, which helps LOCC with distinguishing what is useful as predictors of outcome. While the LOCC score is correlated with the cox regression coefficients and −log(p-values) (Sup Fig S4A), we believe LOCC score provides more information particularly for prognosis. One example of the difference between the cox regression p-value and LOCC score is the rankings of *KIF20A*, *TTK*, and *TPX20.* Although the three gene expressions are strongly correlated (Sup Fig 4B-D), LOCC scoring values very significant cutoffs while cox regression favors having a more consistent HR (Table 1). While the LOCC score does favor significant cutoffs, it is not as singular as using only the most significant cutoff as required by some established methods (Budczies, et al., 2012; Nagy, et al., 2021). Instead as a hybrid score, LOCC is more holistic and can function without adjustment, though future studies will be needed to evaluate LOCC score utility in full gene expression prognosis analysis.

There are still many limitations and issues to be solved in the field of bioinformatics with regard to categorization of continuous variables. Focusing on cancer and gene expression, LOCC can help with ranking variables and visualizing the relevance of the variable and outcome, but needs better multi-variable integration. Currently, we are using weighted coefficients from the original study which used LASSO (Least Absolute Shrinkage Selector Operator) regression. While many models like LASSO use single weight approaches, new methods may be needed for optimizing LOCC score and prognosis (Friedman, et al., 2010). This is true because the LASSO regression discards variables that may be active predictors especially if predictors are correlated or if the number of predictors is not significantly larger than the number of samples (Friedman, et al., 2010; Tibshirani, et al., 2012). Many highly prognostic genes are strongly correlated: for example, *KIF20A*, *TTK*, and *TPX20* in hepatocellular carcinoma all have correlation coefficients above 0.85 (Sup Fig S4B-D). LASSO regression may discard one or more of these variables during its regression process. Furthermore, it is clear from LOCC visualization that gene expression may be significant throughout the entire spectrum or only for a certain range. Thus, a continuous weighted score is appropriate for some variables but unsuitable for others. Biologically, this is understandable because a certain level of protein or gene expression may be required for or representative of activation while lower levels do not signify any meaningful biological difference. Thus, a better method of integrating these variables is anticipated and remains a challenge in research (McDermott, et al., 2013).

One valuable aspect of LOCC is by visualizing the selection process and hazard risks, it is similar to showing one’s work in mathematics. The audience can easily see what cutoffs are selected and how they were selected while also seeing how the validation worked or did not work. Although there are still many calculations performed behind the computer, it is easier to check other’s work by seeing if the LOCC graphs match. Also when validation does not work as intended, there can be visualization of what went wrong as such no peak or a weaker than anticipated signal. Therefore, LOCC can be used to supplement existing and future studies involving prognosis by showing the complete landscape of the predictor or biomarker and increase the confidence of the audience.

### 4.2 Limitations

LOCC and other cutoff selection methods are prone to overfitting and the p-values selected at the best cutoff should be not be interpreted as typical p-values. Nevertheless, we believe LOCC and its scoring can help alleviate simple overfitting by single point cutoff because it visualizes the entire distribution of p-values and hazard ratios. Additionally, the LOCC score incorporates a range component which helps combat overfitting at a specific cutoff. One aspect of LOCC that is both a limitation and benefit is the dependency on the p value, meaning that large datasets can perform better with significance. However, it is also an advantage to incorporate p values to assist in our understanding and selection of prognostic predictors. Ultimately, the best way to verify a predictor is through external validation with another dataset.

## 5. Conclusion

LOCC is a visualization tool that can appropriately display the vast information of continuous variables with regard to survival or prognosis. LOCC can help with cutoff selection and analyze the extent in which a variable is significantly associated with an outcome. The LOCC score can help identify variables that best associate with survival throughout the entire range and rank predictors that allows for discrimination of good and poor predictors. LOCC visualization and score can be used instead of ROC curves for prognosis as it provides more relevant information regarding outcomes.

## Data Availability

Gene expression and mutation data was accessed from cBioportal.com for TCGA LIHC and https://dcc.icgc.org/projects/LIRI-JP for LIKI-JP. Code for LOCC on R will be available at the time of publication.

## Author Contributions

George Luo: Conceptualization, Methodology, Software, Validation, Visualization, Writing – original draft, Writing – review & editing. John J Letterio: Supervision, Validation, Writing – review & editing.

## Acknowledgments

G.L is supported by NIH MSTP training grant 5T32GM007250. J.J.L. is supported by the Jane and Lee Seidman Chair in Pediatric Cancer Innovation. The graphical abstract was created with Biorender.

## Supporting Information

Supplemental Figure S1. Detailed labeling of LOCC graphs

Supplemental Figure S2. Visual and Mathematical Comparison of LOCC and ROC curve

Supplemental Figure S3. Analysis of 12-gene original RISK score using LOCC Supplemental Figure S4. Correlations between LOCC score and cox regression

