## Supplemental Figures for "LOCC: a novel visualization and scoring of cutoffs for continuous variables"

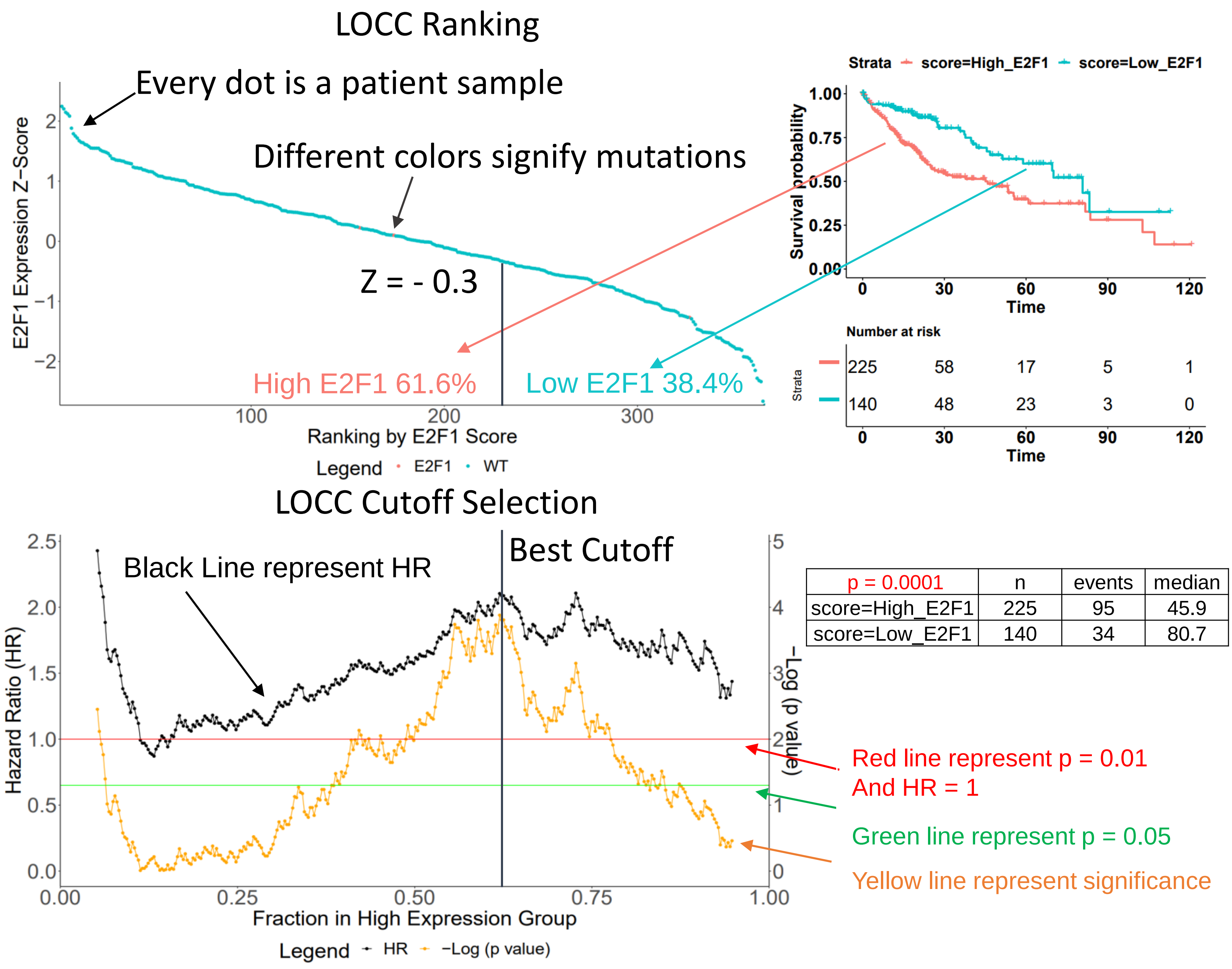


| p = 0.0001 | n | events | median |
|---|---|---|---|
| score=High_E2F1 | 225 | 95 | 45.9 |
| score=Low_E2F1 | 140 | 34 | 80.7 |

All tumor samples are ranked by E2F1 expression
Both graphs' samples and x-axis are lined up

Fig S1.

A B

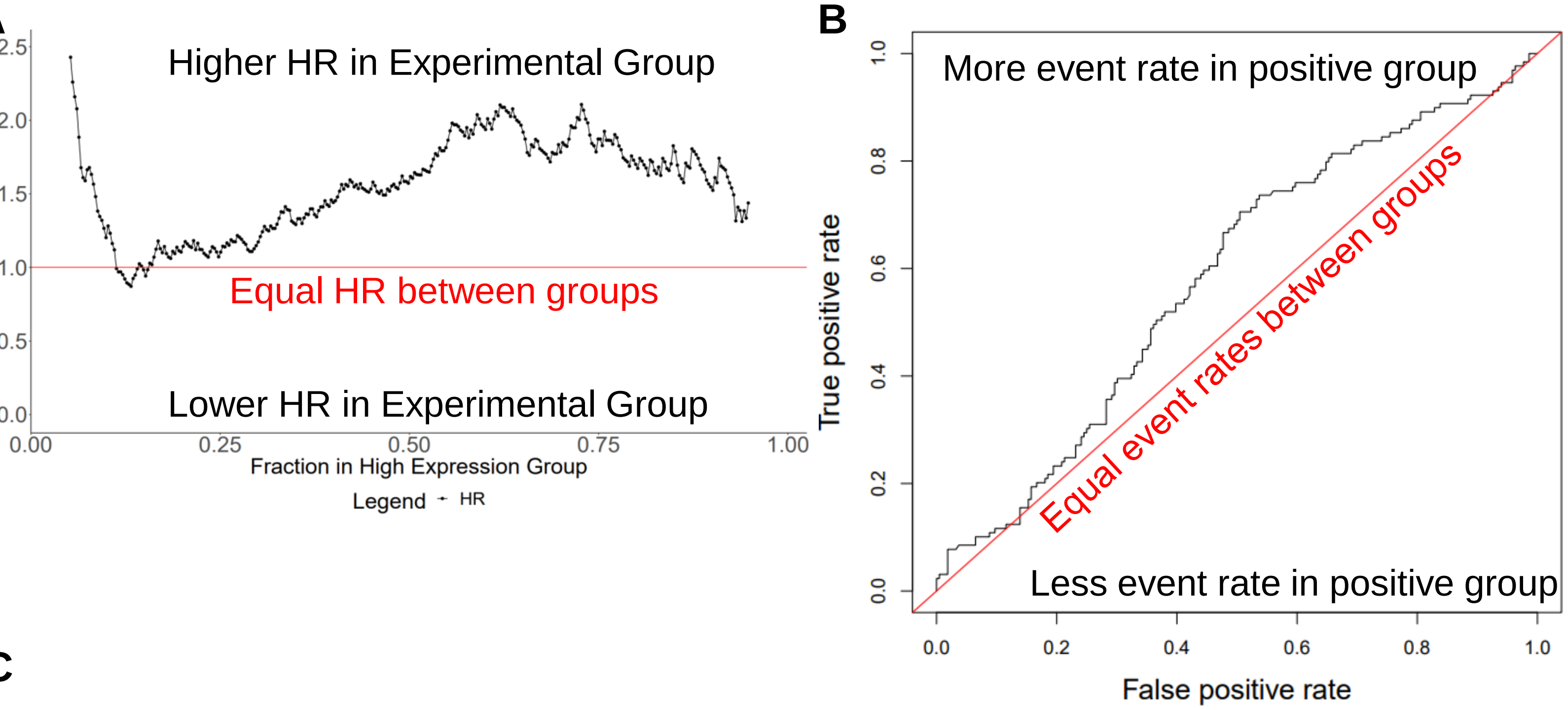


C

| Test | Death | No Death |
|---|---|---|
| Positive | True positive (TP) | False Positive (FP) |
| Negative | False Negative (FN) | True Negative (TN) |

Sensitivity $= \frac{TP}{TP+FN}$ Specificity $= \frac{TN}{TN+FP}$

True Positive Rate (TPR) = Sensitivity

False Positive Rate (FPR) = 1 - Specificity

Positive Predictive Value (PPV) $= \frac{TP}{TP+FP}$

Negative Predictive Value (NPV) $= \frac{TN}{TN+FN}$

Hazard ratio $= \frac{\frac{TP}{TP+FP}}{\frac{FN}{FN+TN}} = \frac{PPV}{1-NPV}$

D

If hazard ratio = 1, then

(1) $\frac{\frac{TP}{TP+FP}}{\frac{FN}{FN+TN}} = 1$

(2) $\frac{TP}{TP+FP} = \frac{FN}{FN+TN}$

(3) TP(FN+TN) = FN(TP+FP)
(4) TP*FN + TP*TN = FN*TP+FN*FP
(5) TP*TN = FN*FP

E

If Sensitivity = 1 – Specificity (TPR=FPR), then

(1) $\frac{TP}{TP+FN} = (1- \frac{TN}{TN+FP}) = \frac{FP}{TN+FP}$

(2) TP(TN+FP) = FP(TP+FN)
(3) TP*TN + TP*FP = FP*TP + FP*FN
(4) TP*TN = FP*FN

Fig S2.

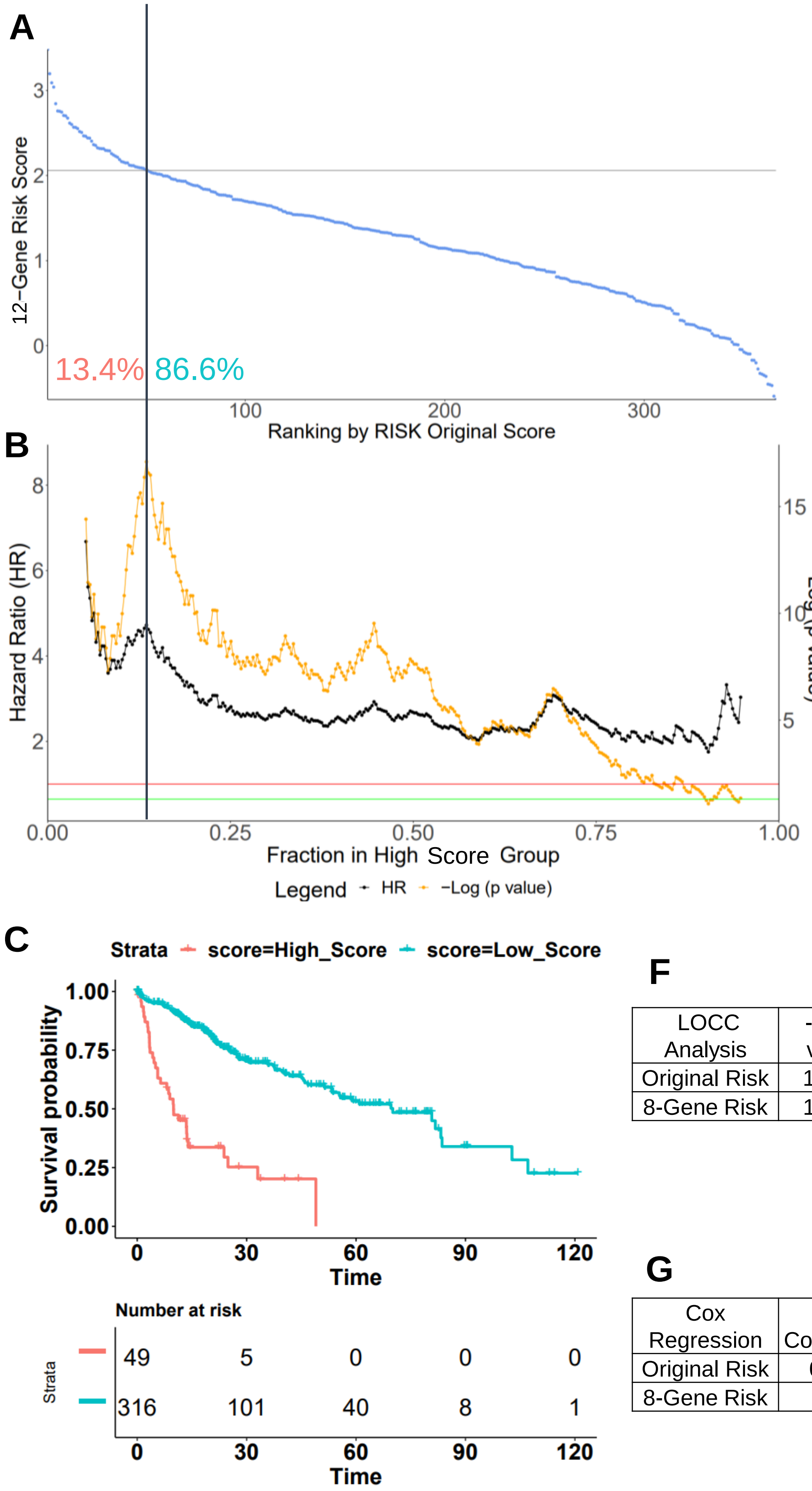


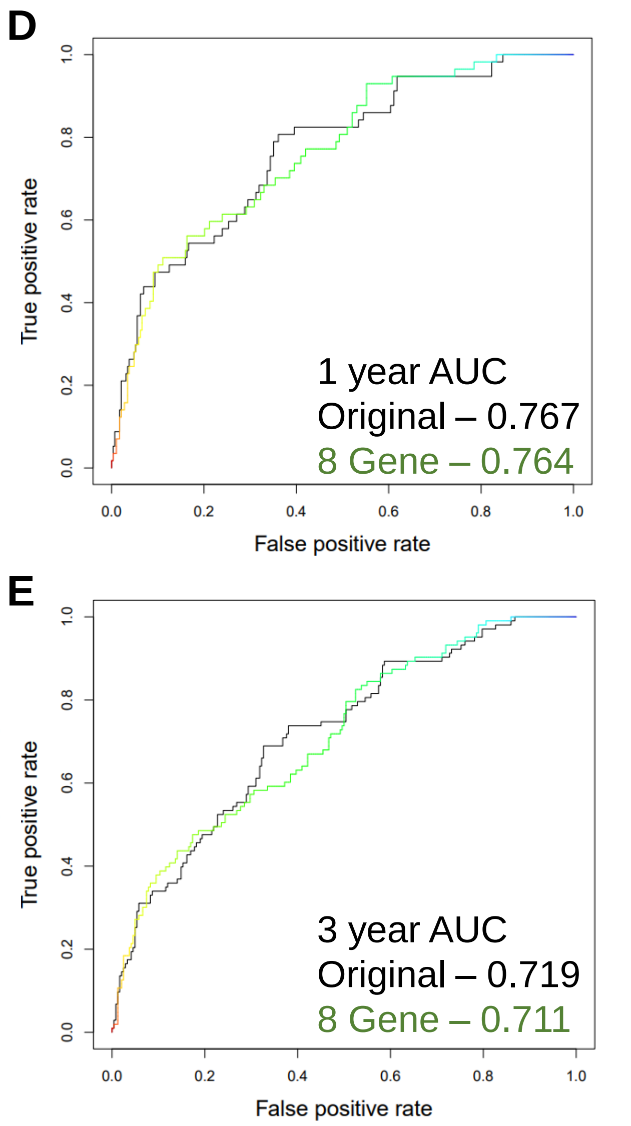


**F**

| LOCC Analysis | -log (p value) | Percentage (<0.01) | Highest HR | Lowest HR | LOCC Score |
|---|---|---|---|---|---|
| Original Risk | 17.093 | 0.837 | 4.723 | 1.942 | 67.6 |
| 8-Gene Risk | 17.008 | 0.871 | 4.971 | 1.972 | 73.6 |

**G**

| Cox Regression | β Coefficient | Exp (β) | Concordance | Log rank test (p value) |
|---|---|---|---|---|
| Original Risk | 0.893 | 2.44 | 0.699 | 1.0E-11 |
| 8-Gene Risk | 1.04 | 2.82 | 0.696 | 2.0E-10 |

Fig S3.

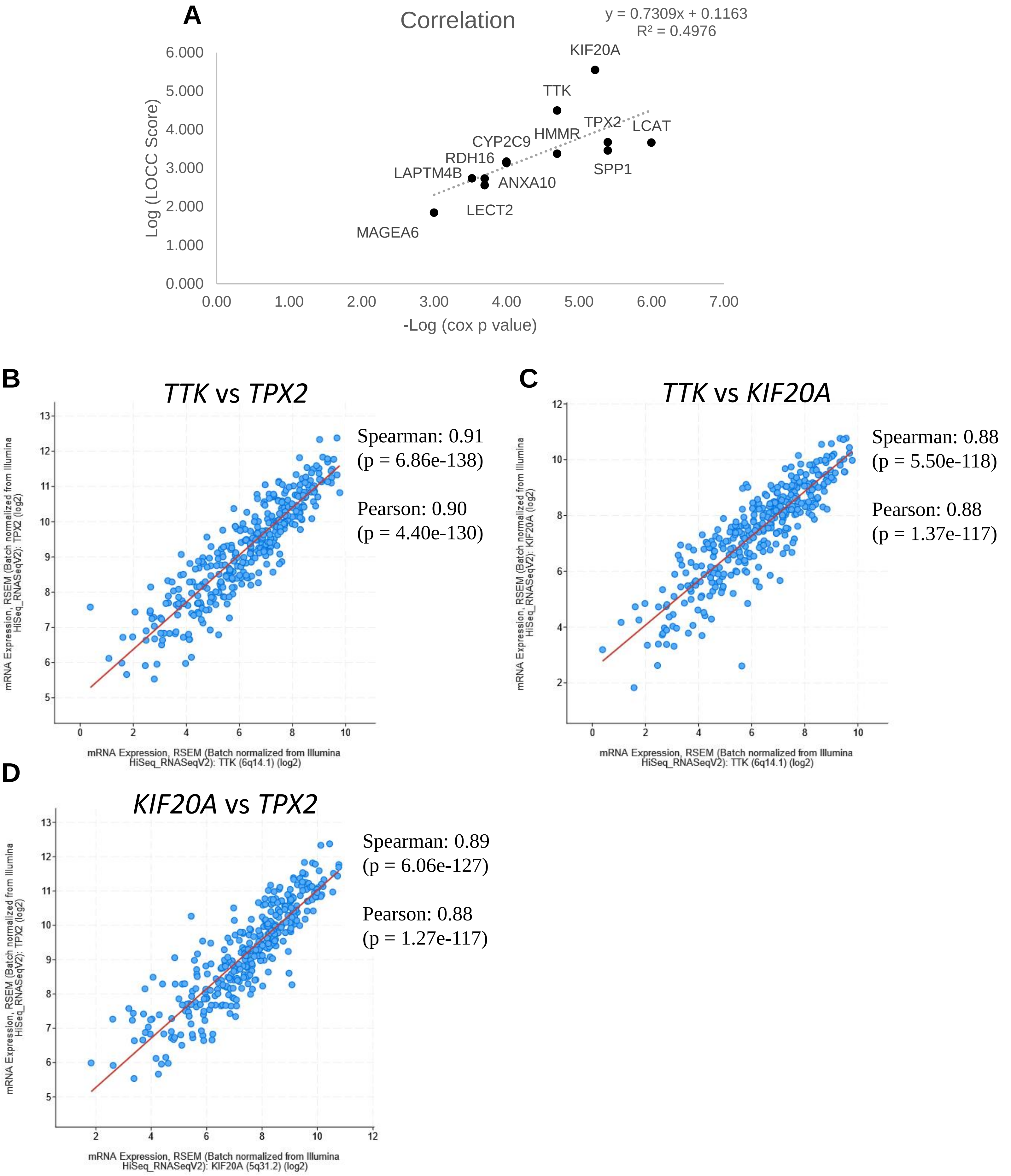
A
Correlation
y = 0.7309x + 0.1163
R² = 0.4976
Log (LOCC Score)
-Log (cox p value)
KIF20A
TTK
TPX2
LCAT
HMMR
CYP2C9
RDH16
LAPTM4B
ANXA10
SPP1
LECT2
MAGEA6
B
TTK vs TPX2
Spearman: 0.91
(p = 6.86e-138)
Pearson: 0.90
(p = 4.40e-130)
mRNA Expression, RSEM (Batch normalized from Illumina HiSeq_RNASeqV2): TTK (6q14.1) (log2)
mRNA Expression, RSEM (Batch normalized from Illumina HiSeq_RNASeqV2): TPX2 (log2)
C
TTK vs KIF20A
Spearman: 0.88
(p = 5.50e-118)
Pearson: 0.88
(p = 1.37e-117)
mRNA Expression, RSEM (Batch normalized from Illumina HiSeq_RNASeqV2): TTK (6q14.1) (log2)
mRNA Expression, RSEM (Batch normalized from Illumina HiSeq_RNASeqV2): KIF20A (log2)
D
KIF20A vs TPX2
Spearman: 0.89
(p = 6.06e-127)
Pearson: 0.88
(p = 1.27e-117)
mRNA Expression, RSEM (Batch normalized from Illumina HiSeq_RNASeqV2): KIF20A (5q31.2) (log2)
mRNA Expression, RSEM (Batch normalized from Illumina HiSeq_RNASeqV2): TPX2 (log2)

Fig S4.
