## Supplemental Figure Legends for "LOCC: a novel visualization and scoring of cutoffs for continuous variables"

**Supplemental Figure 1. Detailed labeling of LOCC graphs.** The LOCC ranking is a graph that plots individual samples values and their respective ranking across all samples. The samples are color coded according to mutation status. The x-axis of ranking directly corresponds to the x-axis of the LOCC cutoff selection graph such that each sample are the same. Therefore, a vertical line from the cutoff selection matches the sample expression and ranking on the LOCC ranking. The cutoff selection has many lines labeled and color coded for ease of visualization. Ideally, the LOCC ranking and cutoff selection are set up so that the hazard ratio is above the red line (HR = 1 as it is easier to visualize HR above 1 than HR below 1. The best cutoff is usually the one with the most significance which is the lowest p-value, visually shown by a peak in the yellow line. However, exceptions can occur if one peak is very close to the edge while comparable peaks exist closer to the middle of the graph. After the cutoff is selected, a corresponding Kaplan-Meier curve is graphed to match the cutoff. The two groups from the Kaplan-Meier curve correspond to the two groups separated by the cutoff on the LOCC ranking. Theoretically, a Kaplan-Meier graph can be plotted for every point on the LOCC graphs which is why there is a large amount of information contained in LOCC.

**Supplemental Figure 2. Visual and mathematical comparison of LOCC and ROC curve. (A)** The HR line of the LOCC graph for *E2F1* expression in TCGA hepatocellular carcinoma was plotted in black. The red line represent a HR = 1. If the HR is above the red line, there are more a higher risk of death associated with the experimental group whereas if the black line is below the red line, there is a lower risk of death associated with the experimental group. (**B**) A ROC curve was plotted for the *E2F1* expression in TCGA hepatocellular carcinoma. A red line is used to show where the true positive rate (TPR) equals the false positive rate (FPR). This line is also referred to as a random classifier because it cannot differentiate true or false positives. Above the red line is where there is an increased rate of events in the test group while being below the red line is a decreased rate of events in the test group. (**C**) A table shows the groups classified by the predictor and the outcome. Statistical equations and abbreviations are listed. (**D**) Equations are calculated under the assumption HR is 1 to calculate the relationship between groups. (**E**) Equations are calculated under the assumption TPR = FPR to calculate the relationship between groups.

**Supplemental Figure 2. Analysis of 12-gene original RISK score using LOCC (A)** The 12-gene original RISK score was ordered in descending order and plotted against the ranking of the samples. A horizontal line is graphed at 2.05 to separate patients into high and low risk groups using ideal LOCC cutoff. **(B)** The LOCC cutoff selection was graphed for the 12-gene RISK score using TCGA hepatocellular data. The most significant cutoff was chosen to separate patients into two groups. (**C**) A Kaplan-Meier overall survival curve is plotted at the validation cutoff to separate patients into high or low risk score. (**D**- **E**) Time-dependent ROC curve of risk score model for 1- and 3-year overall survival predictions were plotted for the original and modified 8-gene RISK scores. (**F**) A full LOCC score analysis is shown comparing the original 12-gene RISK score and the modified 8-gene RISK score. Patients’ survival times are expressed in months. P-values are calculated using log-rank test. HR are calculated using Cox proportional hazard regression.

**Supplemental Figure 3. Correlations between LOCC score and cox regression (A)** A scatterplot is plotted between the Log_2_(LOCC Score) and -Log_10_ (cox p-value). There is a strong correlation (r = 0.705) between the two markers. **B-D.** Scatterplots of gene expression are plotted between the following genes: (**B**) *TTK* vs *TPX2*, (**C**) *TTK* vs *KIF20A*, (**D**) *KIF20A* vs *TPX2*.
